# Self-Motion Perception Influences Postural Sway More than Environmental Motion Perception

**DOI:** 10.1101/2025.05.02.651511

**Authors:** Eric R. Anson, Kyle Critelli, Edward Chen, Jeffrey P. Staab, Mark G. Carpenter, Benjamin Crane

## Abstract

Motion of the visual field can alter postural sway and cause illusions of self-motion. The relative perceptual sensitivity of self-motion versus visual field motion induced by virtual reality (VR) stimulation and whether observable sway differs based on perceptual task is unknown. Methods to quantify sway perception while concurrently measuring sway do not exist. We measured head sway and motion perception (self or world) in healthy adults who stood with feet together wearing a VR headset while experiencing adaptive staircases of virtual sinusoidal pitch rotation about the ankle axis. In separate conditions of randomly ordered blocks, subjects were asked to indicate (yes/no) if the room moved (regardless of perceived postural sway) or if their postural sway increased (regardless of perceived room motion). Head sway area was measured by tracking movement of the VR headset. Yes/No responses were fit with psychometric curves to determine points of subjective equality (PSEs) for room motion and postural sway. PSEs were compared between conditions. Effects of motion perception (binary responses) on head sway area before, during, and after visual stimulation were examined. The mean PSE for room motion (0.42 degrees) was significantly lower than for postural sway (2.02 degrees) [t(1,18) = 4.4714, p = 0.00029]. Head sway area was significantly larger during (z = 11.53, p < 0.001) and after (z = 5.09, p < 0.001) visual stimulation only when participants perceived increased postural sway. Nearly 5-fold greater amplitudes of oscillating VR visual stimuli were required to induce perceptions of altered self-versus visual field motion. Observed head sway during visual motion was only linked to perceptual responses when participants focused internally on self-motion, not externally on room motion.

## Introduction

Quantification of individuals’ perception of self-motion and its relationship to observed (measured) motion remains challenging in clinical and research settings because self-motion perception is confounded by multiple variables. These include ambiguities inherent in natural motion stimuli (e.g., self-motion versus environmental motion as source of visual flow) [1,2], human elements that specifically affect motion perception (e.g., attentional bias directed toward self-motion versus environmental motion and potential induction of illusory motion sensations of vection) [3,4], as well as human elements that generally affect reporting of subjective experiences such as the tendency to amplify ratings of initial exposures to stimuli (elevation bias) or react to the presence of observers (reaction bias) [5–8].

Further, these variables may interact with one another. Although it is well known that ambiguous visual motion may be interpreted as either self-motion (egocentric) or environmental motion (allocentric) [9–11], it is unclear how these interpretations are affected by the focus of attention or context of the task at hand. Studies that manipulated attention found that directing attention internally versus externally could exacerbate the ambiguity of visual motion stimuli and produce mixed effects on postural control [12–14]. Subjects may not have been able to maintain an internal focus on postural sway while purposefully viewing perceptible (external) visual motion stimuli.

Visual motion has long been known to modulate postural sway [15–17], and induces changes in sway from baseline that may be either perceptible (supra-threshold) or imperceptible (sub-threshold). Curiously, the amplitude of sway induced by visual motion stimuli attenuates as stimulus amplitude increases [15,18]; perhaps associated with a shift from using visual stimuli to align the body (i.e., egocentric frame of reference at lower stimulus amplitudes) to interpreting visual stimuli as environmental movement apart from the body (allocentric frame of reference at higher stimulus amplitudes). Thus, even though visually-induced postural sway would be more perceptible at larger amplitudes of visual stimuli [19], it is unclear whether the window between the perceptual threshold and postural sway attenuation is too narrow for adequate perceptual detection of visually induced change in natural postural sway. Despite this unknown, prior work on postural sensory thresholds suggests that suprathreshold sway would be perceived [20].

Visual motion stimuli also induce vection [21–23], with the illusory sensations of motion perceived as either egocentric or allocentric movements [9]. Thus, standing individuals experiencing vection during optic flow may interpret it as either a change in postural sway or environmental motion around them. The strength of vection increases with complexity of the visual motion stimuli (e.g., many objects moving in the same direction simultaneously versus few objects moving in different directions) [24,25]. An immersive virtual reality (VR) scene may induce strong vection, a potential confound for postural experiments in VR.

Unfortunately, the current state of the art for measuring motion perception relies on self-report questionnaires and visual analogue scales that were not designed to account for these confounding variables and their interactions [26–28]. VR, which has been used increasingly in clinical care and research to address problems of balance and postural control [29–33], may offer a tool to quantify self-motion perception while controlling some of these confounds. For example, VR stimuli that are presented in a single plane should elicit less vection than more complex stimuli that often occur in the natural world. Embedding two-alternative forced choice paradigms (2AFC) within VR protocols would allow manipulation of attention and measurement of the effects of egocentric versus allocentric focus on perception of self-motion and environmental motion. Cybersickness from either vection or unintended increased postural sway can lead to feelings of anxiety and/or instability which may be influenced by the degree of presence in the VR environment [34,35]. It is unclear whether self-reported measures of fear/anxiety, or perceived stability will differ when the same VR scenario is used to probe perception of either increased postural sway (internally focused condition) or room motion (externally focused condition).

Here we utilized a VR headset to present visual stimuli of a virtual room moving sinusoidally in the pitch plane to healthy volunteers standing in a relaxed posture. We utilized a standard 2AFC paradigm in which participants had to respond “yes” or “no” to perceptions of an increase in postural sway (internally focused condition) or room motion (externally focused condition) to determine the amplitude of visual stimulus amplitude corresponding to the point of subjective equality (PSE) for perceived postural sway increase and perceived room motion postural sway. We hypothesized that the PSE for detecting an increase in visually-induced postural sway would be larger than the PSE for detecting room motion. Although multisensory integration improves perceptual detection of self-motion [36], the perceptual discrimination tasks investigated here have different levels of complexity. In the case of room motion detection, the comparison is motion to no motion. However, in the case of sway increase, the comparison is baseline sway to greater sway. Change in ongoing postural sway may require greater cognitive processing compared to visual motion detection and is susceptible to both sensory and process noise [37]. We also hypothesized that perceiving increased postural sway would correspond to observable increases in postural sway (i.e., observable movement rather than vection). We hypothesized that self-reported measures of fear and anxiety would not differ between the conditions, but that self-reported stability may be lower when subjects were internally focused on postural sway perception.

## Materials and Methods

### Subjects

Twenty-five healthy adults (13 women, 12 men) mean age 33.8 (± 20.4), provided written informed consent and participated in this study, approved by the Institutional Review Board at the University of Rochester Medical Center. A power analysis based on preliminary data and sway data [38], suggested a sample of 10-15 was needed to detect changes in perception or sway area. We recruited 25 as the effects for visual perturbation may differ from vestibular perturbation, and not all subjects perform ideally in a 2AFC paradigm. At the time of data collection, all persons at the University of Rochester Medical Center were required to wear facemasks at all times. Participants were compensated $15 per hour.

### Experimental Setup Apparatus

The HTC VIVE VR System (HTC Corporation, Taoyuan, Taiwan) was used for the study. The head-mounted display (HMD) presented stereoscopic images at a resolution of 1080x1200 pixels per eye, resulting in a 110° field of view with an average refresh rate of 90Hz. Custom Python scripts were run via VR software (Vizard, WorldVis, Inc, Santa Barbara, CA USA) on a Desktop computer (Dell Technologies, Round Rock, TX USA) to control the motion of the virtual environment. HMD position was captured at 90Hz as a proxy for postural sway [39,40]. Anterior-posterior and mediolateral sway trajectories were used to calculate sway area for the head [41].

### Protocol

After consenting, each participant provided demographic data including age, gender. Participants were familiarized with the 2AFC paradigm with a brief demonstration protocol that included virtual visual motion and a prompt to respond by clicking either a virtual “yes” or “no” button using the HTC hand-held controller.

Participants were instructed to stand upright and relaxed with their feet together, avoiding a stiff posture. Participants were tested in two blocked conditions (visual motion detection or perceived sway increase) in random order determined by a coin flip. Each condition consisted of 50 trials (each trial consisted of 16 seconds of VR exposure followed by a 2AFC question). During the first and last 3 second periods of each trial, there was no imposed room motion. During the middle 10 seconds, two cycles of 0.2 Hz visual motion were presented at varied amplitudes based on responses to the 2AFC paradigm [42], see Figure 1. A randomly interleaved 2AFC adaptive staircase design was used with 25 trials per staircase [43], see Figure 1.

**Figure 1.**
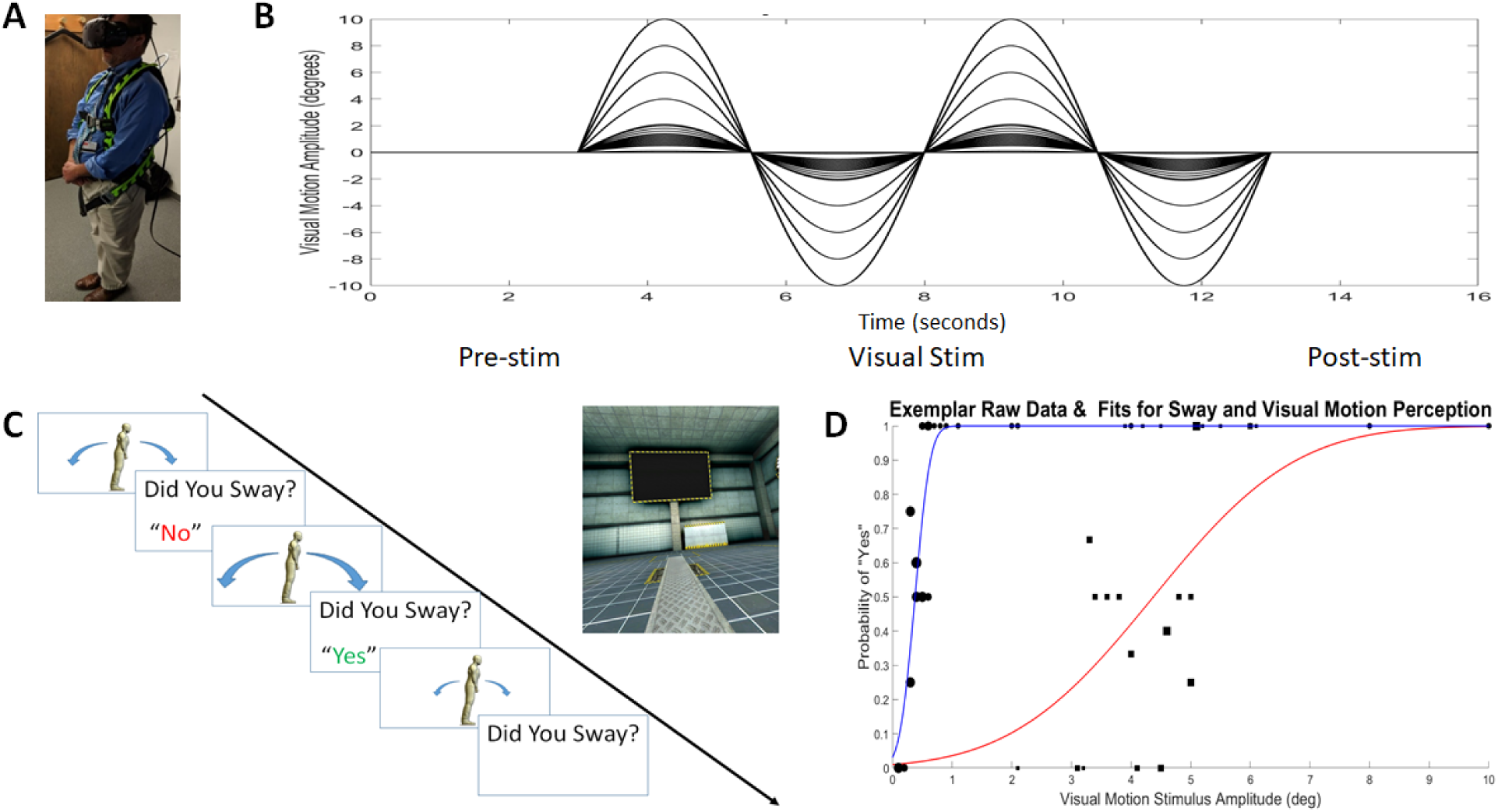
Methods for the experiment. A) Example of subject (Author EA who consented to including this image) standing position wearing the VR headset. B) Exemplar visual motion stimuli: 3 seconds of no motion, 10 seconds of 0.2 Hz sinusoids, 3 seconds of no motion. Minimum amplitude 0.1 degree, maximum amplitude 10 degrees. C) Example of the response paradigm. “Yes” responses lead to smaller stimuli and “no” responses to larger stimuli at the next presentation. D) Example binary response data (circles = perceived room motion, squares = perceived postural sway) and psychometric fits. The x-value at the y=0.5 level corresponds to the PSE.

For both staircases, the maximum visual amplitude was 10 degrees and the minimum visual amplitude was 0.1 degrees. The initial staircase direction was determined randomly. Three consecutive steps in the same direction on a staircase or a staircase reversal resulted in switching staircases. The next step(s) must be for the other staircase. Additionally, if all 25 trials were completed for a particular staircase, the remaining trials must all be from the other staircase. The maximum and minimum step sizes were specified as 2 degrees and 0.1 degrees. The initial step size for each staircase was set to the maximum step size. For the descending staircase, the next step size was doubled following a “yes” response and halved following a “no” response. For the ascending staircase, the next step size was doubled following a “no” response and halved following a “yes” response. If the next step size was larger (smaller) than the specified maximum (minimum) step size, the step size was re-specified as the maximum (minimum) step size. The visual motion amplitude was adjusted based on prior responses to converge on the point of subjective equality (PSE) which corresponds to the visual motion amplitude where the subject was equally likely to identify neighboring visual amplitude values as the same (i.e., present or absent). This method focused the majority of stimuli near the PSE [44]. The inter-trial interval varied based only on time to respond to the 2AFC question, and participants were also instructed to use the 2AFC question time to shift their weight or move slightly to minimize fatigue. Head motion was not tracked during this time.

After completing both sets of staircases, self-reports were collected with subjects quantifying fear, state anxiety, perceived postural stability, VR immersiveness, and realism of the virtual environment on a 0-100 point scale using a slider bar that appeared in the VR scene [45,46].

### Data Analysis

Data are available at University of Rochester Figshare website: https://doi.org/10.60593/ur.d.28886831

All data were analyzed offline using custom MATLAB scripts (MathWorks, Inc, Natick, MA, USA) and Stata version 14 (StataCorp LLC, College Station, TX USA). As a first pass, the ascending and descending staircases were examined for convergence based on overlapping histograms to ensure appropriate task performance. The binary data (no = 0, yes = 1) were bootstrapped with 100 replications and fit to a psychometric curve. The mean of the bootstrapped psychometric fits was calculated as the PSE (in degrees) for each condition and participant.

To test our hypothesis that PSE for detecting increased sway would be greater than for detecting room motion, we used an ANOVA to compare PSEs for detecting visual motion and increased postural sway while controlling for age and sex.

To test our hypothesis that perceived increased postural sway would correspond to observed increased postural sway we conducted two sets of analyses. First, we used a linear mixed model analysis to compare sway area across time (before, during, and after the visual perturbation) and perceptual response type (“yes/no”) while controlling for age and sex to determine if there was an interaction between perturbation timing and perceptual response. We expected that “yes” perceptual responses when internally focused would correspond to measured increase in postural sway compared to “no” responses.

Second, we examined the transfer function between visual motion and postural sway to restrict the analysis to postural sway attributable to the visual perturbation excluding aspects of postural sway incoherent with the visual stimulus [47]. Power spectral densities (PSD) and cross spectral densities (CSD) were calculated for the visual motion and anterior-posterior head displacement during the visual stimulation using MATLAB functions *pwelch* and *cpsd*. Sway gain to vision was calculated as the absolute value of the transfer function defined as CSD_vision-sway_/PSD_vision_ at the visual motion frequency 0.2 Hz [48].

Sway gain to vision had a non-linear relationship with visual stimulus amplitude. Therefore, an exploratory multivariable hierarchical fractional polynomial regression was performed regressing sway gain on amplitude (visual motion amplitude) and response (yes/no) to determine the correct polynomial power for amplitude and determine whether both independent variables were needed in the model. The exploratory multivariable hierarchical fractional polynomial regression determined a coefficient of -0.5 was appropriate for visual motion amplitude and perceptual response was also important to the model with a coefficient of 1. To determine whether visually driven sway differed according to perceptual response (yes/no) we regressed sway gain on amplitude (visual motion amplitude) and perceptual response using the following regression model:

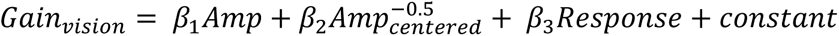

*AMP_centered_* was used to reduce collinearity with *Amp,* which was included to improve overall model fit. The only statistical comparison used from this regression was whether the relationship between postural gain to vision and amplitude was dependent on perceptual response (𝛽_3_ in the above equation).

To test the hypothesis that self-reported fear and anxiety would not differ, but that self-reported stability would be lower when subjects were internally focused on postural sway perception, we performed separate paired t-tests compared self-reports (fear, anxiety, stability, immersiveness, reality) between conditions. Alpha was specified for the PSE ANOVA analysis and multiple regression effects as α = 0.05, a Bonferroni corrected α = 0.003 for the mixed model sway area analysis, and a Bonferroni corrected α = 0.01 for condition effects on the paired t-tests for self-reports.

## Results

Five subjects did not demonstrate staircase convergence and were excluded from further analyses. The average age of the 20 analyzed subjects was 37.6 (21.2) years with 10 women and 10 men.

The average PSE for perceived room motion (0.44 (0.2) degrees) was significantly lower than the average PSE for perceived postural sway increase (2.15 (1.57) degrees) [F(1,39) = 22.4, p = 0.0001]. Age and sex were not significant covariates in the ANOVA model comparing PSE between conditions.

There was a significant main effect of visual stimulation periods (before, during, after) on sway area for both the internal (Χ^2^ (2,18) = 339.05, P < 0.001) and external (Χ^2^ (2,18) = 504.57, P < 0.001) perceptual focus conditions, see Figure 2. Observed postural sway area significantly increased during the visual perturbation for both perceived postural sway (z = 20.61, p < 0.001) and room motion (z = 15.52, p < 0.001) conditions compared to the pre-stimulus sway area. There was a significant interaction between perceptual response to postural sway increase (yes/no) and visual stimulation period (before, during, after) on sway area (Χ^2^ (2,18) = 68.37, P < 0.001) only for the internal perceptual focus. Observed postural sway area was significantly larger during (z = 11.53, p < 0.001) and after (z = 5.09, p < 0.001) the visual perturbation for “yes” compared to “no” responses only for the internally focused (perceived postural sway) condition, see Figure 2A. Age and sex did not have a significant effect on sway area.

**Figure 2.**
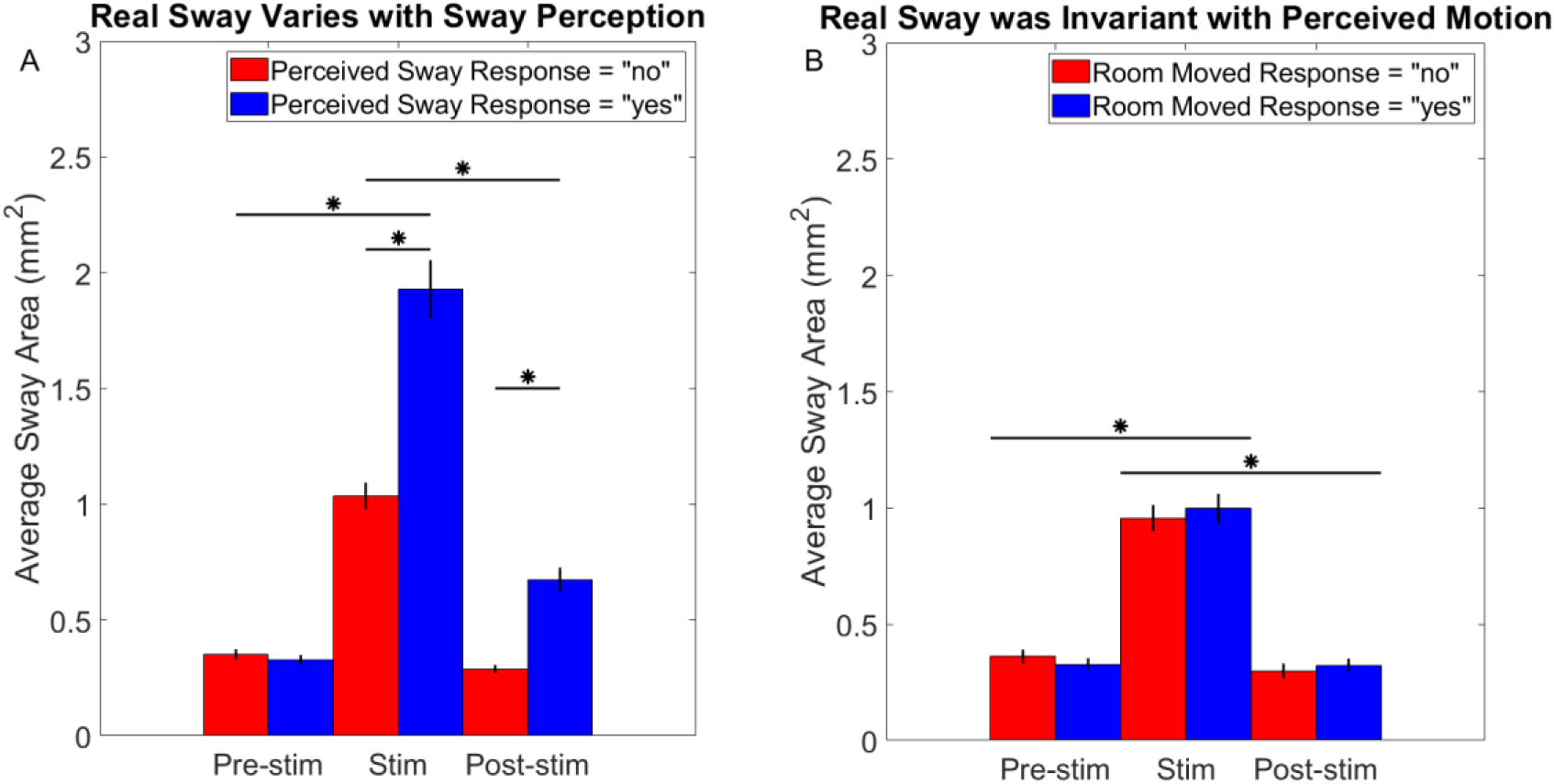
Observed sway area as a function of perceptual response (yes = blue, no = red). Observed sway area significantly increased during the visual stimulation (Stim) for both conditions: A) Perceived postural sway, and B) Perceived room motion. Only in the perceived postural sway condition was there significantly greater average observed head sway area when subjects responded yes. There was a significant after effect in the post-stimulation period (Post-stim) for the trials with “yes” responses, but only in the perceived postural sway condition.

For the regression analyses, the results of the exploratory multivariable hierarchical fractional polynomial regression are provided in Table 1 for transparency. However, only the subsequent multiple polynomial linear regression (see Table 2) is interpreted. For the perceived postural sway condition, there was a significant visual motion amplitude main effect for gain of observed head sway to vision such that visually driven head sway decreased as visual motion amplitude increased. Age and sex did not have a significant effect on gain of observed head sway to vision.

**Table 1.** Exploratory multinomial fractional polynomial regression results. Initial model comparisons (linear versus fractional polynomial) and determination of final model. All predictors were considered significant in the model.

| Variable | Model | Deviance | Deviance Difference | Significance | Powers |
| --- | --- | --- | --- | --- | --- |
| AMP | Linear vs. Fractional Polynomial | 702.24 | 466.27 | $p < 0.001$ | 1 |
| | Fractional Polynomial | 238.09 | 2.11 | $p = 0.350$ | -0.5 |
|  | Final Model | 238.09 |  |  | -.05 |
| Independent Variables | Regression Coefficients | Standard Error | T score | Significance | 95% Confidence |
| AMP<sup>-.5</sup> | 0.44 | 0.01 | 30.7 | $p < 0.001$ * | [0.41 – 0.47] |
| Response (y/n) | 0.10 | 0.02 | 5.54 | $p < 0.001$ * | [0.06 – 0.14] |
| Constant | 0.12 | 0.01 | 8.36 | $p < 0.001$ * | [0.09 – 0.15] |

**Table 2.** Polynomial regression results. The overall polynomial regression was significant (F(3,9996) = 317.62, p < 0.001) and the R-squared = 0.49. Untransformed visual motion amplitude was included to improve the fit of model predictions but was not overall a significant contributor to the model.

| Dependent Variable | Coefficient | Standard Error | Significance | 95% Confidence |
| --- | --- | --- | --- | --- |
| AMP | -0.0039 | 0.0047 | $p = 0.404$ | [-0.013– 0.0053] |
| AMP<sup>-.5</sup> | 0.43 | 0.018 | $p < 0.001^*$ | [0.39 – 0.46] |
| Response (y/n) | 0.01 | 0.018 | $p < 0.001^*$ | [0.066 – 0.14] |
| Age | 0.0001 | 0.0004 | $p = 0.799$ | [-0.0007 – 0.0009] |
| Sex | 0.02 | 0.018 | $p = 0.249$ | [-0.01 – 0.05] |
| Constant | 0.13 | 0.020 | $p < 0.001^*$ | [0.092 – 0.17] |

For the perceived postural sway condition, there was a significant nonlinear relationship between visual motion amplitude and gain of observed head sway to vision [F(3,996) = 317.62, p < 0.0001]. The overall model explained 49% of the variance in gain of observed head sway to vision. Variance inflation factor (VIF) indicated minimal collinearity (VIF = 1.96) across independent variables. Across amplitudes, the gain of observed head sway to visual perturbation was significantly greater for trials with “yes” responses during the perceived postural sway condition [t(1,996) = 5.6, p < 0.0001], see Figure 3.

**Figure 3.**
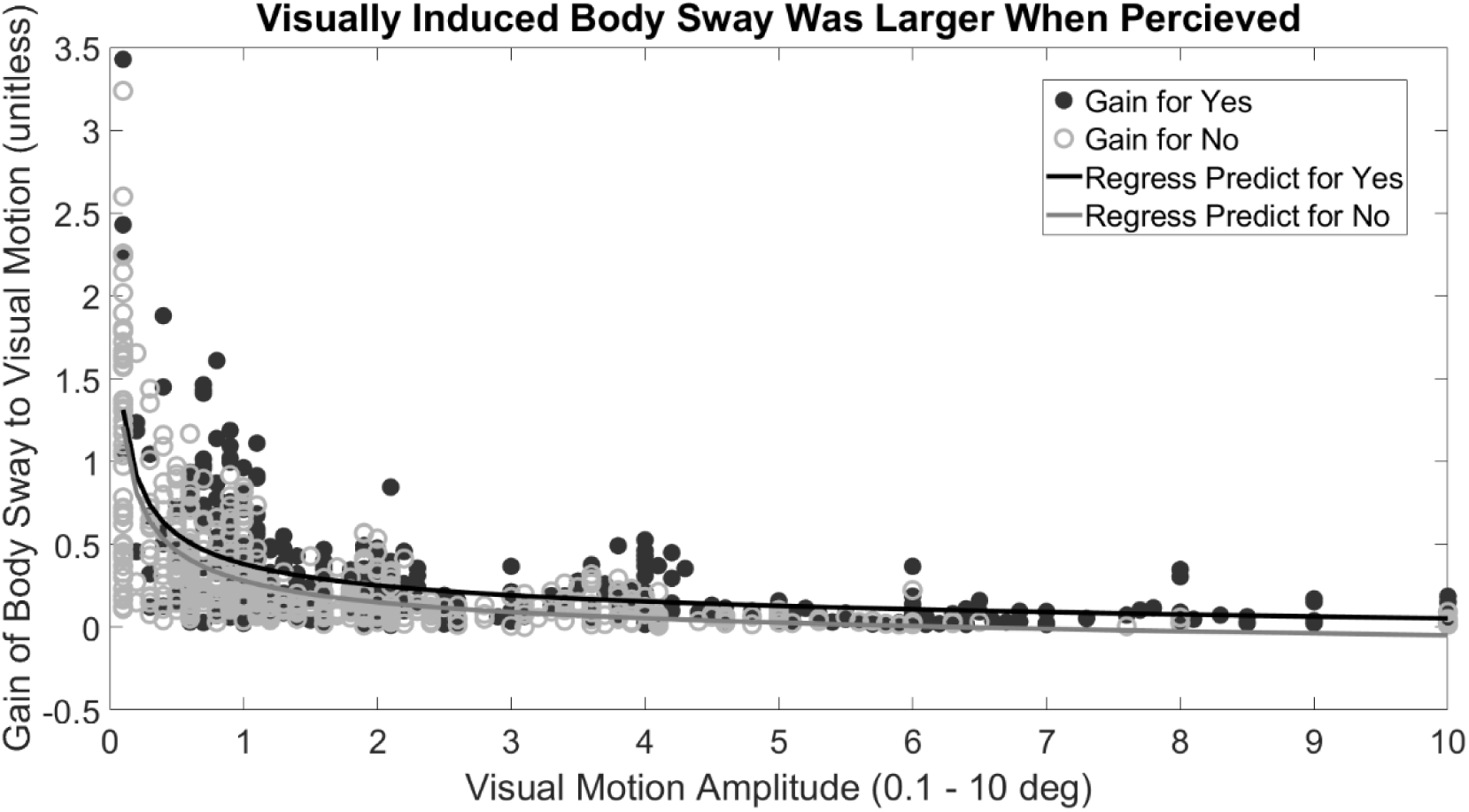
Gain to vision as a function of visual motion amplitude and perceptual response for the perceived postural sway condition. Filled circles represent individual steps on the perceptual staircases for all individuals (grey = “no,” and black = “yes” responses). Solid lines represent polynomial fits (grey = “no,” and black = “yes” responses) derived from the polynomial regression (see Table 4). Gain of postural sway to visual motion was significantly greater when subjects responded “yes” to perceiving an increase in sway during the self-motion perception condition.

The average values for the self-reported measures (fear, anxiety, stability, immersiveness, realism) did not differ between conditions (p’s > 0.2), presented in Table 3.

**Table 3.** Average (SD) subjective ratings for both conditions (perceived room motion and perceived postural sway increase)

| Rating | Perceived Room Motion | Perceived Postural sway Increase | p value |
| --- | --- | --- | --- |
| Fear | 8.4 (14.2) | 6.6 (13.2) | p = 0.2145 |
| Anxious | 13.4 (22.6) | 15.6 (21.6) | p = 0.5064 |
| Stability | 62.3 (27.9) | 66.5 (27.3) | p = 0.4788 |
| Immersion | 64 (28) | 60.9 (28.7) | p = 0.3635 |
| Real | 57.6 (27.3) | 60 (27.8) | p = 0.4197 |

## Discussion

In the current study, subjects were able to focus internally on postural sway despite perceivable apparent room motion and externally on a moving visual stimulus despite increased postural sway, which indicates that perception of self-motion and environmental motion are separable, even when the movements occur together.

The PSE for perceived visual motion in this study was larger than previously reported thresholds for anterior-posterior visual motion detection (0.06-0.12 degrees) [20]. This discrepancy can be attributed to several methodological differences. First, and potentially the most important difference, our subjects were not immobilized. Perceptual judgements of stimulus magnitude are known to be sensitive to sensory noise. Self-motion is known to increase the just noticeable difference for visual motion detection [49], thus natural body sway may account for most of the discrepancy. Second, VR headsets are known to restrict the peripheral field of view (approximately 110 degrees). Peripheral vision is important for motion detection, and reduced peripheral vision may reduce sensitivity for visual motion detection or alter attribution from allocentric to egocentric motion perception [9]. Taken together, unrestricted sway and reduced field of view as well as perceptual testing methods likely account for the larger than expected PSE for visual motion detection in VR.

As expected, observed postural sway increased significantly from baseline during the visual motion period whether perceptual focus was internally or externally directed. Interestingly, observed postural sway was significantly greater when subjects reported a perceived increase in postural sway compared to trials when they responded that their sway had not increased. There was also an after-effect, such that sway area remained larger during the post-stim period (Figure 2A), only during trials when subjects reported a perceived increase in postural sway for the internally focused “perceived postural sway increased” condition. This suggests that the subjects’ perceptual responses were aligned with an actual increase in postural sway rather than a vection perception (illusion of self-motion).

Increasing visual velocity leads to sway attenuation attributed to sensory reweighting [42,48,50,51], and our current results with varied motion amplitude during the perturbation period are consistent with this effect (see Figure 3). In our paradigm, increased visual perturbation amplitude at the fixed frequency of 0.2 Hz corresponds to increased visual motion velocity. Our data are interesting in that the frequency response for head sway to vision followed a similar overall pattern as head sway area during the stimulation period such that perceived visually driven sway responses were larger.

When perceptual focus was internally directed, head sway gain in response to visual motion was higher when subjects indicated “yes” their sway had increased. Others recently reported that visual weighting was similar between a control condition and a condition of focusing internally on sway [52]. Our observation that differences in visually driven sway response when perceptual focus was internally directed depended on whether increased postural sway was perceived. Although both studies involved internally or externally focused balance attention, a key difference in the current study was the perceptual sway estimation. In our case, the change in head sway gain appears to be mediated by the additional processing to judge change in sway rather than observe ongoing sway. Future studies are needed to clarify the role of different types of perceptual focus on sensory weighting.

It is not clear why an after-effect was observed for the “perceived sway increased” trials, see Figure 2A. One interpretation is that larger magnitude visually driven sway responses may have taken longer to return to baseline. Alternatively, sway not directly associated with the visual perturbation (for example postural sway at frequencies other than 0.2 Hz) may also explain the after-effect. Postural sway in the presence of visual perturbation contains both congruent (visually driven) and incongruent (not associated with visual perturbation signals) body movements [47]. This after-effect was not observed when subjects were externally focused, highlighting the task dependency. A recent study examining postural sway effects in VR reported that sinusoidal stimuli only elicited an immediate effect on postural sway [53]. The shorter stimulation paradigm and inclusion of different perceptual loci (internal versus external) may contribute to the discordance between the present results and that recent report [53] . The internal perceptual focus of the current study may have facilitated longer entrainment to the repeated sinusoidal visual perturbation in the current study. When perceptual focus was directed externally, our results (lack of after-effect) are more consistent with the prior study [53].

Others have demonstrated that postural sway increased more when visual perturbations in VR were expected [54]. Although all participants in the current study were aware that the virtual room would move on some trials, the adaptive nature of the interleaved staircases should have reduced the subjects’ expectation that the room would move for any given trial [43], especially for the externally focused “perceived room motion” paradigm. As the staircases began to converge for the “perceived postural sway” paradigm, subjects may have more consistently expected the room to move. In this study, subjects were not queried regarding their expectations for room motion and future studies should explore this as a potential for the increased sway when responding “yes” during the internally focused paradigm.

In this study, self-reported measures of fear/anxiety, stability, immersiveness and realism did not differ based on perceptual focus (internal vs. external). Others have shown that fear, anxiety, and stability ratings change based on experimental context (i.e., postural threat) [55,56]. In our study, both conditions presented visual motion leading to an increase in postural sway. Although the observed sway was greater when subjects perceived an increase in sway during the internally focused condition, the post-hoc summative self-report of stability did not reflect that condition difference. It is not clear whether the lack of condition difference for self-reported stability ratings represents a limitation of the self-reflective question (each condition lasted approximately 20 minutes) or whether this composite represents a regression to the average perceived postural sway. Further, it is possible that self-reported stability ratings are not based on the observed sway since psychological ratings do not always change consistently with observed sway across repeated exposures [57]. Using the post-hoc self-reported fear and anxiety as proxies for arousal [57–59], suggests that the VR experience was only mildly arousing and perceptual focus likely did not impact arousal. Future studies should measure trial by trial effects using physiological measures of arousal and instantaneous stability ratings.

## Limitations

It is possible that sway increased in response to the perception (illusion) that their sway had increased. This seems unlikely, since visually coupled head motion was present during trials when sway increase was not perceived, albeit at smaller magnitudes. Future studies should explore the contributions of baseline sway to self-motion perception as well as the ability to identify individuals with anxiety-related or function vestibular disorders for whom diagnostic biomarkers remain elusive.

## Conclusion

Individuals appear able to interpret similar visual scene motion differently based on task demands. Greater amplitudes of VR visual oscillations were required to induce perceptions of altered self-compared to virtual visual field motion. Observable head sway related to accurate motion perceptions only when internally focused (self-motion), not when externally focused (room motion). Visually driven postural sway was influenced by perception of change in ongoing sway, indicating that not all postural sway induced by visual sensory feedback reaches the level of perceptual awareness. This technique has the potential to enhance diagnostic capabilities for anxiety-related and functional vestibular disorders such as persistent postural perceptual dizziness or Mal de Debarquement syndrome.

## CRediT author statement

**Eric Anson**: Conceptualization, Investigation, Data curation, Writing-Original draft preparation, Visualization, Methodology, Formal Analysis, Funding acquisition. **Kyle Critelli**: Investigation, Data curation, Methodology, Software, Writing-Reviewing and Editing. **Edward Chen**: Investigation, Software, Writing-Reviewing and Editing. **Jeff Staab**: Supervision, Methodology, Writing-Reviewing and Editing. **Mark Carpenter**: Methodology, Writing-Reviewing and Editing, Supervision. **Benjamin Crane:** Conceptualization, Methodology, Writing-Reviewing and Editing, Resources, Supervision.

## References

[1] A. Piedimonte, A.J. Woods, A. Chatterjee, Disambiguating ambiguous motion perception: what are the cues?, Front Psychol 6 (2015) 137879. 10.3389/FPSYG.2015.00902/BIBTEX.

[2] H.C. Longuet-Higgins, Visual motion ambiguity, Vision Res 26 (1986) 181–183. 10.1016/0042-6989(86)90079-9.

[3] G. Wulf, M. Höß, W. Prinz, Instructions for motor learning: differential effects of internal versus external focus of attention., J Mot Behav 30 (1998) 169–79. 10.1080/00222899809601334.

[4] K.A. Becker, C.J. Hung, Attentional focus influences sample entropy in a balancing task, Hum Mov Sci 72 (2020). 10.1016/j.humov.2020.102631.

[5] N. Hendrix, N. Maisel, J. Everson, V. Patel, A. Bazemore, L.S. Rotenstein, A.J. Holmgren, A.H. Krist, J. Adler-Milstein, R.L. Phillips, Impact of response bias in three surveys on primary care providers’ experiences with electronic health records, Journal of the American Medical Informatics Association 31 (2024) 1754–1762. 10.1093/JAMIA/OCAE148.

[6] K.M. Mazor, B.E. Clauser, T. Field, R.A. Yood, J.H. Gurwitz, A demonstration of the impact of response bias on the results of patient satisfaction surveys, Health Serv Res 37 (2002) 1403– 1417. 10.1111/1475-6773.11194.

[7] J.A. Pulido, L.H. Barrero, S.E. Mathiassen, J.T. Dennerlein, Correctness of Self-Reported Task Durations: A Systematic Review, Ann Work Expo Health 62 (2017) 1–16. 10.1093/ANNWEH/WXX094.

[8] P.E. Shrout, G. Stadler, S.P. Lane, M. Joy McClure, G.L. Jackson, F.D. Clavél, M. Iida, M.E.J. Gleason, J.H. Xu, N. Bolger, Initial elevation bias in subjective reports, Proc Natl Acad Sci U S A 115 (2018) E15–E23. 10.1073/PNAS.1712277115/ASSET/B5DC3D51-9A55-4272-9BF1-C34B1F2F72B5/ASSETS/GRAPHIC/PNAS.1712277115FIG02.JPEG.

[9] T. Brandt, J. Dichgans, E. Koenig, Differential effects of central versus peripheral vision on egocentric and exocentric motion perception, Exp Brain Res 16 (1973) 476–491. 10.1007/BF00234474.

[10] D. Alais, R. Keys, F.A.J. Verstraten, C.L.E. Paffen, Vestibular and active self-motion signals drive visual perception in binocular rivalry, IScience 24 (2021) 103417. 10.1016/J.ISCI.2021.103417.

[11] H.C. Longuet-Higgins, Visual motion ambiguity, Vision Res 26 (1986) 181–183. 10.1016/0042-6989(86)90079-9.

[12] N. Richer, D. Saunders, N. Polskaia, Y. Lajoie, The effects of attentional focus and cognitive tasks on postural sway may be the result of automaticity, Gait Posture 54 (2017) 45–49. 10.1016/J.GAITPOST.2017.02.022.

[13] T. Kim, J. Jimenez-Diaz, J. Chen, The effect of attentional focus in balancing tasks: A systematic review with meta-analysis, Journal of Human Sport and Exercise 12 (2017) 463–479. 10.14198/JHSE.2017.122.22.

[14] T.J. Ellmers, G. Machado, T.W.L. Wong, F. Zhu, A.M. Williams, W.R. Young, A validation of neural co-activation as a measure of attentional focus in a postural task, Gait Posture 50 (2016) 229–231. 10.1016/j.gaitpost.2016.09.001.

[15] J. Jeka, T. Kiemel, R. Creath, F. Horak, R. Peterka, Controlling human upright posture: velocity information is more accurate than position or acceleration., J Neurophysiol 92 (2004) 2368–79.

[16] J.R. Lishman, D.N. Lee, The autonomy of visual kinaesthesis, Perception 2 (1973) 287–294. 10.1068/p020287.

[17] C.W. Hoppes, P.J. Sparto, S.L. Whitney, J.M. Furman, T.J. Huppert, Functional near-infrared spectroscopy during optic flow with and without fixation, PLoS One 13 (2018) e0193710. 10.1371/journal.pone.0193710.

[18] R.J. Peterka, Sensorimotor integration in human postural control, J Neurophysiol 88 (2002) 1097– 1118.

[19] D.M. Merfeld, Signal detection theory and vestibular thresholds: I. Basic theory and practical considerations, in: Exp Brain Res, Springer Verlag, 2011: pp. 389–405. 10.1007/s00221-011-2557-7.

[20] R. Fitzpatrick, D.I. McCloskey, Proprioceptive, visual and vestibular thresholds for the perception of sway during standing in humans., J Physiol 478 (1994) 173–186. 10.1113/jphysiol.1994.sp020240.

[21] M. Moroz, I. Garzorz, E. Folmer, P. MacNeilage, Sensitivity to visual speed modulation in head-mounted displays depends on fixation, Displays 58 (2019) 12–19. 10.1016/j.displa.2018.09.001.

[22] A. Nehrujee, L. Vasanthan, A. Lepcha, S. Balasubramanian, A Smartphone-based gaming system for vestibular rehabilitation: A usability study, J Vestib Res 29 (2019). 10.3233/VES-190660.

[23] M.T. Robert, L. Ballaz, M. Lemay, The effect of viewing a virtual environment through a head-mounted display on balance, Gait Posture 48 (2016) 261–266. 10.1016/J.GAITPOST.2016.06.010.

[24] B. Keshavarz, H. Hecht, Axis rotation and visually induced motion sickness: The role of combined roll, pitch, and yaw motion, Aviat Space Environ Med 82 (2011) 1023–1029. 10.3357/ASEM.3078.2011.

[25] B. Keshavarz, A.E. Philipp-Muller, W. Hemmerich, B.E. Riecke, J.L. Campos, The effect of visual motion stimulus characteristics on vection and visually induced motion sickness, Displays 58 (2019) 71–81. 10.1016/j.displa.2018.07.005.

[26] M. Schieppati, E. Tacchini, A. Nardone, J. Tarantola, S. Corna, Subjective perception of body sway., J Neurol Neurosurg Psychiatry 66 (1999) 313–22. 10.1136/JNNP.66.3.313.

[27] P. Castro, D. Kaski, M. Schieppati, M. Furman, Q. Arshad, A. Bronstein, Subjective stability perception is related to postural anxiety in older subjects, Gait Posture 68 (2019) 538–544. 10.1016/J.GAITPOST.2018.12.043.

[28] E. Anson, S. Studenski, P.J. Sparto, Y. Agrawal, Community-dwelling adults with a history of falling report lower perceived postural stability during a foam eyes closed test than non-fallers, Exp Brain Res 237 (2019) 769–776. 10.1007/s00221-018-5458-1.

[29] A. V. Lubetzky, D. Harel, H. Darmanin, K. Perlin, Assessment via the oculus of visual “Weighting” and “Reweighting” in young adults, Motor Control 21 (2017) 468–482. 10.1123/mc.2016-0045.

[30] M. Moroz, I. Garzorz, E. Folmer, P. MacNeilage, Sensitivity to visual speed modulation in head-mounted displays depends on fixation, Displays 58 (2019) 12–19. 10.1016/j.displa.2018.09.001.

[31] A. Nehrujee, L. Vasanthan, A. Lepcha, S. Balasubramanian, A Smartphone-based gaming system for vestibular rehabilitation: A usability study, J Vestib Res 29 (2019). 10.3233/VES-190660.

[32] M.T. Robert, L. Ballaz, M. Lemay, The effect of viewing a virtual environment through a head-mounted display on balance, Gait Posture 48 (2016) 261–266. 10.1016/J.GAITPOST.2016.06.010.

[33] F. Delgado, C. Der Ananian, The Use of Virtual Reality Through Head-Mounted Display on Balance and Gait in Older Adults: A Scoping Review, Games Health J 10 (2021) 2–12. 10.1089/G4H.2019.0159.

[34] B. Arcioni, S. Palmisano, D. Apthorp, J. Kim, Postural stability predicts the likelihood of cybersickness in active HMD-based virtual reality, Displays 58 (2019) 3–11. 10.1016/j.displa.2018.07.001.

[35] D. Gromer, M. Reinke, I. Christner, P. Pauli, Causal Interactive Links Between Presence and Fear in Virtual Reality Height Exposure, Front Psychol 10 (2019) 141. 10.3389/fpsyg.2019.00141.

[36] M.W. Greenlee, S.M. Frank, M. Kaliuzhna, O. Blanke, F. Bremmer, J. Churan, L.F. Cuturi, P.R. MacNeilage, A.T. Smith, Multisensory Integration in Self Motion Perception, Multisens Res 29 (2016). 10.1163/22134808-00002527.

[37] S. Nouri, F. Karmali, Variability in the Vestibulo-Ocular Reflex and Vestibular Perception, Neuroscience 393 (2018) 350–365. 10.1016/j.neuroscience.2018.08.025.

[38] R.C. Fitzpatrick, S.R.D. Watson, Passive motion reduces vestibular balance and perceptual responses, Journal of Physiology 593 (2015) 2389–2398. 10.1113/JP270334.

[39] B. Sylcott, K. Williams, M. Hinderaker, C.-C. Lin, Comparison of HTC Vive^TM^ Virtual Reality Headset Position Measures to Center of Pressure Measures, Proceedings of the Human Factors and Ergonomics Society Annual Meeting 63 (2019) 2333–2336. 10.1177/1071181319631344.

[40] A. V. Lubetzky, Z. Wang, T. Krasovsky, Head mounted displays for capturing head kinematics in postural tasks, J Biomech 86 (2019) 175–182. 10.1016/J.JBIOMECH.2019.02.004.

[41] F. Danion, M.L. Latash, eds., Motor Control Theories, Experiments, and Applications, Oxford University Press, Inc., New York, 2011.

[42] R.J. Peterka, Sensorimotor integration in human postural control, J Neurophysiol 88 (2002) 1097– 1118.

[43] T.N. Cornsweet, The staircrase-method in psychophysics., Am J Psychol 75 (1962) 485–491. 10.2307/1419876.

[44] R.E. Roditi, B.T. Crane, Suprathreshold asymmetries in human motion perception., Exp Brain Res 219 (2012) 369–79. 10.1007/s00221-012-3099-3.

[45] T.W. Cleworth, B.C. Horslen, M.G. Carpenter, Influence of real and virtual heights on standing balance., Gait Posture 36 (2012) 172–6. 10.1016/j.gaitpost.2012.02.010.

[46] A.L. Adkin, J.S. Frank, M.G. Carpenter, G.W. Peysar, Fear of falling modifies anticipatory postural control., Exp Brain Res 143 (2002) 160–70. 10.1007/s00221-001-0974-8.

[47] E. Anson (2025). Dataset for vision and sway motion perception balance study. University of Rochester. Dataset. 10.60593/ur.d.28886831

[48] E. Anson, P. Agada, T. Kiemel, Y. Ivanenko, F. Lacquaniti, J. Jeka, Visual control of trunk translation and orientation during locomotion, Exp Brain Res 232 (2014) 1941–1951.

[49] T. Kiemel, A.J. Elahi, J.J. Jeka, Identification of the plant for upright stance in humans: multiple movement patterns from a single neural strategy., J Neurophysiol 100 (2008) 3394–406. 10.1152/jn.01272.2007.

[50] A.H. Wertheim, G. Reymond, Neural noise distorts perceived motion: the special case of the freezing illusion and the Pavard and Berthoz effect., Exp Brain Res 180 (2007) 569–76. 10.1007/s00221-007-0887-2.

[51] R.J. Peterka, M.S. Benolken, Role of somatosensory and vestibular cues in attenuating visually induced human postural sway, Exp Brain Res 105 (1995) 101–110. 10.1007/BF00242186.

[52] J. Jeka, T. Kiemel, R. Creath, F. Horak, R. Peterka, Controlling human upright posture: velocity information is more accurate than position or acceleration., J Neurophysiol 92 (2004) 2368–79.

[53] L. Ma, P.J. Marshall, W.G. Wright, The impact of external and internal focus of attention on visual dependence and EEG alpha oscillations during postural control, J Neuroeng Rehabil 19 (2022). 10.1186/S12984-022-01059-7.

[54] J. Ketterer, S. Ringhof, D. Gehring, A. Gollhofer, Sinusoidal Optic Flow Perturbations Reduce Transient but Not Continuous Postural Stability: A Virtual Reality-Based Study, Front Physiol 13 (2022) 803185. 10.3389/FPHYS.2022.803185/BIBTEX.

[55] H. Chander, S.N.K.K. Arachchige, C.M. Hill, A.J. Turner, S. Deb, A. Shojaei, C. Hudson, A.C. Knight, D.W. Carruth, Virtual-reality-induced visual perturbations impact postural control system behavior, Behavioral Sciences 9 (2019). 10.3390/bs9110113.

[56] T.W. Cleworth, B.C. Horslen, M.G. Carpenter, Influence of real and virtual heights on standing balance., Gait Posture 36 (2012) 172–6. 10.1016/j.gaitpost.2012.02.010.

[57] T.J. Ellmers, E.C. Kal, W.R. Young, Consciously processing balance leads to distorted perceptions of instability in older adults, J Neurol 268 (2021) 1374–1384. 10.1007/s00415-020-10288-6.

[58] M. Zaback, A.L. Adkin, M.G. Carpenter, Adaptation of emotional state and standing balance parameters following repeated exposure to height-induced postural threat, Sci Rep 9 (2019) 12449. 10.1038/s41598-019-48722-z.

[59] T.W. Cleworth, M.G. Carpenter, Postural threat influences conscious perception of postural sway, Neurosci Lett 620 (2016) 127–131. 10.1016/j.neulet.2016.03.032.

[60] A.L. Adkin, M.G. Carpenter, New Insights on Emotional Contributions to Human Postural Control, Front Neurol 9 (2018) 789. 10.3389/fneur.2018.00789.

